# Downregulation of RyR and NCX in the neonatal rat ventricular myocyte modulates cytosolic [Ca^2+^]

**DOI:** 10.1101/2021.10.20.465171

**Authors:** Esteban Vazquez-Hidalgo, Xian Zhang, David Torres Barba, Paul Paolini, Parag Katira

## Abstract

Calcium (Ca^2+^) is necessary for cardiac muscle contraction. RyR, NCX, and SERCA are key regulatory protein channels for cytosolic Ca^2+^ in cardiac myocytes. Expression levels of these proteins are a function of development, with protein expression shifting toward the adult phenotype over time. We investigated how downregulation by siRNAs of RyR and NCX affected expression levels of complimentary proteins and the corresponding intracellular Ca^2+^ transients. We compared experimentally observed Ca^2+^ transients to those predicted by mathematical models. Experiments show RyR downregulation decreased SERCA and increased NCX protein levels. The associated Ca^2+^ transient had a decreased amplitude, increased time-to-peak, 50%, and 90% Ca^2+^ removal with respect to the control cell. NCX downregulation increased SERCA production without significant changes in RyR expression levels. The corresponding [Ca^2+^] transient had increased amplitude, no change in time-to-peak and 50% Ca^2+^ removal, and increased 90% Ca^2+^ removal with respect to the control cell. Computational models that accurately predict the observed experimental data suggest compensatory changes occurring in the expression levels as well as biochemical activity of the regulatory proteins.

## 1 INTRODUCTION

The heart is a pump that transports necessary blood to the tissues to deliver nutrients, remove metabolic waste, and aid in gas exchange. Tasked with these functions, the heart is the first organ to develop and become functional [1, 2]. Heart beats are detectable in early developmental stages (embryonic day (ED) 22 in humans, ED 9 in rats)[1]. At this stage, the heart is a beating tube with a functioning excitation-contraction coupling (ECC) mechanism [3] that controls electrical signals to mechanical pumping [4], establishing early on the importance of producing controlled cardiac contraction from electrical stimulation and ionic flux balance.

Each heart beat is the result of numerous orchestrated multiscale efforts. Our study is at the cellular and molecular level, focusing on Ca^2+^ flux handling. Ca^2+^ is a necessary ion in heart contraction [5] as the strength of cardiac contraction is regulated by the amplitude and duration of the Ca^2+^ transient [6]. Specifically, we are interested in learning how cytosolic [Ca^2+^] behaves in cultured neonatal rat ventricular myocytes (NRVMs) when short interfering RNAs (siRNA) are used to independently inhibit protein expression of the Na^+^/Ca^2+^ exchanger (NCX1, refered to as NCX) and the cardiac isoform of ryanodine receptors (RyR2, refered to as RyR). Furthermore, we are interested in observing any compensatory changes in protein expression of RyR, NCX, and the sarcoendoplasmic reticulum Ca^2+^-ATPase (SERCA) pump when RyR and NCX are downregulated.

NRVMs are a robust model used to study mechanical and physiological properties [7, 6], develop spatiotemporal computational models of ECC [8, 9], and investigate drug-induced differential gene expression [10]. Similarity between post-natal and diseased adult cardiomyocytes has been used as a rationale for using NRVMs as a model to study certain aspects of cardiovascular disease, such as Ca^2+^ handling and ECC[11]. We acknowledge that NRVMs *in vivo* and *in vitro* are morphologically different [12, 13] from healthy and diseased adult myocytes with the immature myocytes being spheroid while healthy and adult myocytes are rod-like [14]. However, in diseased hearts, transverse tubule (t-tubule) density within the adult myocytes decreases or their position drifts. This drift changes the proximity of the sarcolemmal (SL) L-type Ca^2+^ channel (LTCC) to the sarcoplasmic reticulum (SR) RyR channels [15, 16, 17], essentially leaving “orphaned” RyR channels resulting in altered extracellular and SR Ca^2+^ fluxes [18, 19, 20, 21, 22].

We begin by independent knockdown of RyR and NCX. We measure relative protein expression of RyR, NCX, and SERCA for each knockdown followed by fluorescence intensity measurements of free cytosolic [Ca^2+^] for each knockdown group. Lastly, we use an established computational model calibrated to our experimental conditions to identify underlying dynamics mechanistic implications not explained by differential protein expression.

## 2 MATERIAL AND METHODS

Experiments were carried out in accordance with institutional and federal guidelines. Animal use protocols were approved and supervised by San Diego State University Office of Laboratory Animal Care and the Institutional Animal Care and use Committee.

All procedures were done according to manufacturer’s protocols unless otherwise stated.

### 2.1 Neonatal Rat Ventricular Myocyte Isolation

NRVMs were harvested from litters of 1-day old Sprague-Dawley (*Rattus norvegicus*) hearts. The ventricles were separated from the hearts and placed in air compatible Dulbecco’s Modified Eagle’s Medium (DMEM)(Thermo Fisher, CA). The ventricles were pooled, minced, and enzymatically isolated by seven rounds of trypsin (Invitrogen, CA) at 37°C. After each digestion round, the supernatant was removed and placed in DMEM/F12 (Thermo Fisher, CA) containing 20% fetal bovine serum (FBS) (Thermo Fisher, CA). NRVMs were pre-plated for 2 hours to remove fibroblasts [23].

### 2.2 Gene Silencing

We use downregulation by siRNA as we are interested in altering gene expression in healthy cells vs. using genetically modified cells from animals with congenital abnormalities. Stealth RNAi and scrambled oligo-ribonucleotide duplex (Invitrogen, CA) were used. Reverse transcription of siRNAs were preformed with TransMessenger Transfection Reagent (Qiagen, USA). Fluorescently labeled dsRNA oligomer, BLOCK-iT (Thermo Fisher, CA), optimized delivery rates. Transfections were done prior to NRVM seeding. Gene of interest (GOI) siRNA or control siRNA were incubated at room temperature with Enhancer R and Buffer EC-R (Qiagen, USA) for 20 minutes. TransMessenger Transfection Reagent was added and incubated for an additional 10 minutes at room temperature. Immediately after room temperature incubation, NRVMs with transfection mixtures were placed in DMEM-air at 37°C, 5% CO_2_, for 4-6 hours. Transfection mixtures and DMEM-air were removed and replaced with DMEM/F12 supplemented with 10% FBS, incubated for 48 hours at 37°C and 5% CO_2_.

### 2.3 Western Blots

NRVMs for western blots were pooled and plated in 6-well plates at 0.5 × 10^6^ cells per well from 12 ventricles. NCX and RyR knockdown were performed on separate populations from different litters, each with a control group from the respective group. Protein was extracted 48 hours post transfection with All Prep RNA/Protein Kit (Qiagen, USA). Protein concentrations were determined by Lowry assay (Bio-Rad, CA). 20 *μ*g of total protein was denatured at 70°C for 10 minutes and loaded in 3-8% Tris-Acetate Gel (Invirogen, CA) for NCX and RyR2, running in 1×Tris-glycine SDS Running Buffer (Invitrogen, CA) at 125V for 90 minutes. Proteins were transferred to 0.45 *μ*m pore size nitrocellulose membrane in 1×NuPage Transfer Buffer (Invitrogen, CA). SERCA proteins were loaded in 4-12% Bis-Tris Gels (Invitrogen, CA) running in 1×MOPs SDS running buffer (Invitrogen, CA) at 150V for 90 minutes and transferred to 0.45 *μ*m pore size PVDF membrane (Invitrogen, CA). 0.5% Ponceau-S was used for protein visualization followed by a 5% non-fat milk block buffer for 1 hour at room temperature. Membranes were then transferred to fresh blocking buffer with diluted antibodies (RyR2 1:500, NCX 1:200, SERCA 1:2000, GAPDH 1:5000) (Jackson ImmunoResearch, PA) followed by overnight incubation on a shaker at 4°C. The following day, 3 10-minute TBST washes were performed followed by Western Lightning plus ECL Enhanced Chemiluminescence (Perkin Elmer, MA). Proteins were detected on X-ray film (Fuji, Japan). Protein levels were determined with Image J gel analysis tool kit (NIH, USA). Experiments were performed three times with triplicate samples. Unpaired t-tests were used with population means±SEM. Differences in the means were deemed statistically significant when P< 0.05(^∗^), P< 0.01(^∗∗^), and P< 0.001(^∗∗∗^).

### 2.4 qRT-PCR Analysis

Relative mRNA levels were compared by reverse transcription quantitative polymerase chain reaction (qRT-PCR). Total mRNA was extracted from NRVMs with All Prep RNA/Protein Kit 48 hours post transfection. RNA was amplified in 48-well plates using Eco Real-Time PCR Detection System (Illumina, CA) with KAPA SYBR FAST One-Step qRT-PCR Kits (Kapa Biosystems, MI). NCX, SERCA, and RyR2 RNA samples were analyzed in triplicate using comparative CT value to determine fold change of mRNA expression. Data were normalized by GAPDH (see supplementary tables).

### 2.5 Calcium Transient Measurement

NRVMs for Ca^2+^ measurements were plated at 1×10^6^ cells per well on fibronectin coated coverslips. Ca^2+^ fluorescence intensities were acquired with a dual emission photometry system (Photon Technology International, Lawrenceville, New Jersey) [24]. Briefly, 10*μ*M of Fluo-3AM (Molecular Probes, Inc) in dimethly sulfoxide (DMSO) was added to 10 ml of 10% FBS supplemented with 0.05% pluronic F-127 (Molecular Probes, Inc). Each well was loaded with 1 ml of Fluo-3AM complex and incubated for 20 minutes. After incubation, NRVMs were washed with 1:1 media and the coverslip was placed onto an environmental stage chamber with 1 ml of 1:1 medium. NRVMs were field stimulated at 0.3 Hz for 3 msec duration at 50mV. Fluorescence intensity was measured using Felix computer software (Photon Technology International, Lawrenceville, New Jersey). Due to limitations by the use of non-ratiometric dye, experimental and computational baselines are set to [Ca^2+^] = 0. Cytosolic free [Ca^2+^] was approximated by the pseudoratio

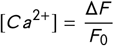

where Δ*F* is the fluorescence intensity and *F*_0_ is the baseline measurement as described in [25] and as performed computationally by Korhonen et al [8].

### 2.6 Mathematical Model

Computational models for ventricular and atrial cardiac systems exist for adult animals. Some of the more recognizable adult ventricular models include rabbit [26], mouse [27], and human [28]. Fewer neonatal models exist. Fewer computational models exist for neonatal animals. There is a neonatal mouse ventricular model [9] and a neonate rat ventricular model [8]. Both models use ordinary differential equations to solve action potential based on ionic concentrations. We employ the mathematical model presented by Korhonen of NRVMs as it closely matches our cell geometry and initial conditions. The model takes into account action potentials based on potassium (K^+^) and sodium (Na^+^) currents as well spatiotemporal cytosolic changes in [Ca^2+^] from the SL and SR. The model provides us with a method to perform a functional analysis of the Ca^2+^ cycling proteins NCX, RyR, and SERCA.

Our focus in this model are the equations related to RyR, NCX, and SERCA. The mathematical model describes the Ca^2+^ efflux from the SR via the RyR channel as

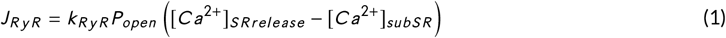

where *k*_*RyR*_ is the RyR scaling factor, [*Ca*^2+^]_*SRrelease*_ is the [Ca^2+^] from the SR, [*Ca*^2+^]_*subSR*_ is the [Ca^2+^] immediately adjacent to the SR membrane, and *P*_*open*_ is the probability function of the channel being in the open state.

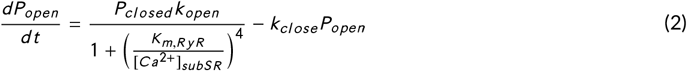

*k*_*open*_ is the rate constant for RyR opening, *k*_*close*_ is the rate constant for RyR closing, and *K*_*m, RyR*_ is the half-saturation for RyR. *P*_*closed*_ is defined as

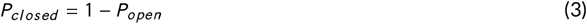

Ca^2+^ uptake into the SR is represented by a Hill-type equation

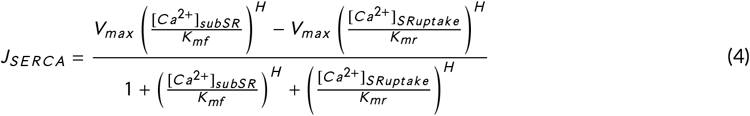

where *V*_*max*_ is interpreted as the SERCA scaling factor, *H* is the Hill coefficient, [Ca^2+^]_*SRuptake*_ is the [Ca^2+^] sequestered into the SR, *K*_*mf*_ is the SERCA forward half-saturation constant, and *K*_*mr*_ is the SERCA reverse half-saturation constant.

Ca^2+^ efflux to the extracellular space by NCX is described as

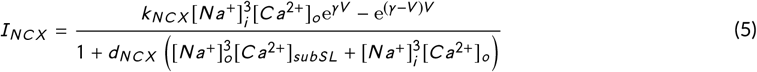

where *k*_*NCX*_ is the NCX scaling factor, *d*_*NCX*_ is the denominator scaling factor, *γ* is the energy barrier parameter, *V* is the total voltage, [Na^+^]_*i*_ is the cytosolic [Na^+^], [Na^+^]_*o*_ is the extracellular [Na^+^], and [Ca^2+^]_*o*_ is the extracelluar [Ca^2+^]. For the full list of model equations, see [8].

Our experimental setup required changes to the model proposed in [8]. At the time of Ca^2+^fluorescence measurements, our cultured NRVMs were larger, 14 *μ*m vs 10*μ*m and our culture media had higher [Ca^2+^], 1.9 mM vs 1.7 mM. The appropriate changes were made in the initial conditions to calibrate the model to obtain a control model for our experimental NRVM (Figure 3, Table 3). Using the observed Ca^2+^ transient from our control NRVM population, we performed minimal RyR, NCX, and SERCA parameters fitting to establish a model that predicted the expected Ca^2+^ response (Figure 3).

## 3 PROTEIN EXPRESSION CHANGES

### 3.1 Downregulation by siRNA

RyR downregulation by siRNA showed a decrease in RyR protein expression by 73% (±9% SEM), an increase in NCX protein expression of 51% (±6% SEM), and a decrease of SERCA expression by 31% (±12% SEM) (Figure 1A, Table 1). The associated Ca^2+^ transients showed decreased amplitude, increased time-to-peak (T_peak_) and increased time to 50% transient decay (T_50_) with respect to the control (Figure 2A).

**TABLE 1.** Western blot expression levels

| Knockdown Condition | RyR Expression | NCX Expression | SERCA Expression |
| --- | --- | --- | --- |
| Control | 1 | 1 | 1 |
| RyR Knockdown | 0.27±0.09** | 1.51±0.06*** | 0.69±.012* |
| NCX Knockdown | 1.25±0.16 | 0.19±0.04*** | 2.04±0.02*** |

Western blot expression levels for each of the knockdown conditions. Shifts in expression levels are reported with respect to the control group. \* $P < 0.05$ , \*\* $P < 0.01$ , \*\*\* $P < 0.001$ .

**TABLE 2.** Ca^2+^ Transient Functional Parameters

| Experimental Condition | Amplitude ($\mu M$) | $T_{peak}(s)$ | $T_{50}(s)$ | $T_{90}(s)$ |
| --- | --- | --- | --- | --- |
| Control | $0.46 \pm 0.03$ | $0.25 \pm 0.02$ | $0.75 \pm 0.02$ | $1.55 \pm 0.09$ |
| RyR Knockdown | $0.33 \pm 0.05^{**}$ | $0.40 \pm 0.01^{**}$ | $1.07 \pm 0.04^{***}$ | $2.25 \pm 0.11^{**}$ |
| NCX Knockdown | $0.59 \pm 0.04^{**}$ | $0.25 \pm 0.01$ | $0.78 \pm 0.02$ | $2.10 \pm 0.03^{**}$ |

Functional parameters of note include $Ca^{2+}$ transient amplitude, time-to-peak ( $T_{peak}$ ), time-to-50% transient decay ( $T_{50}$ ), and time-to-90% decay ( $T_{90}$ ). \* $P < 0.05$ , \*\* $P < 0.01$ , \*\*\* $P < 0.001$ .

**FIGURE 1.**
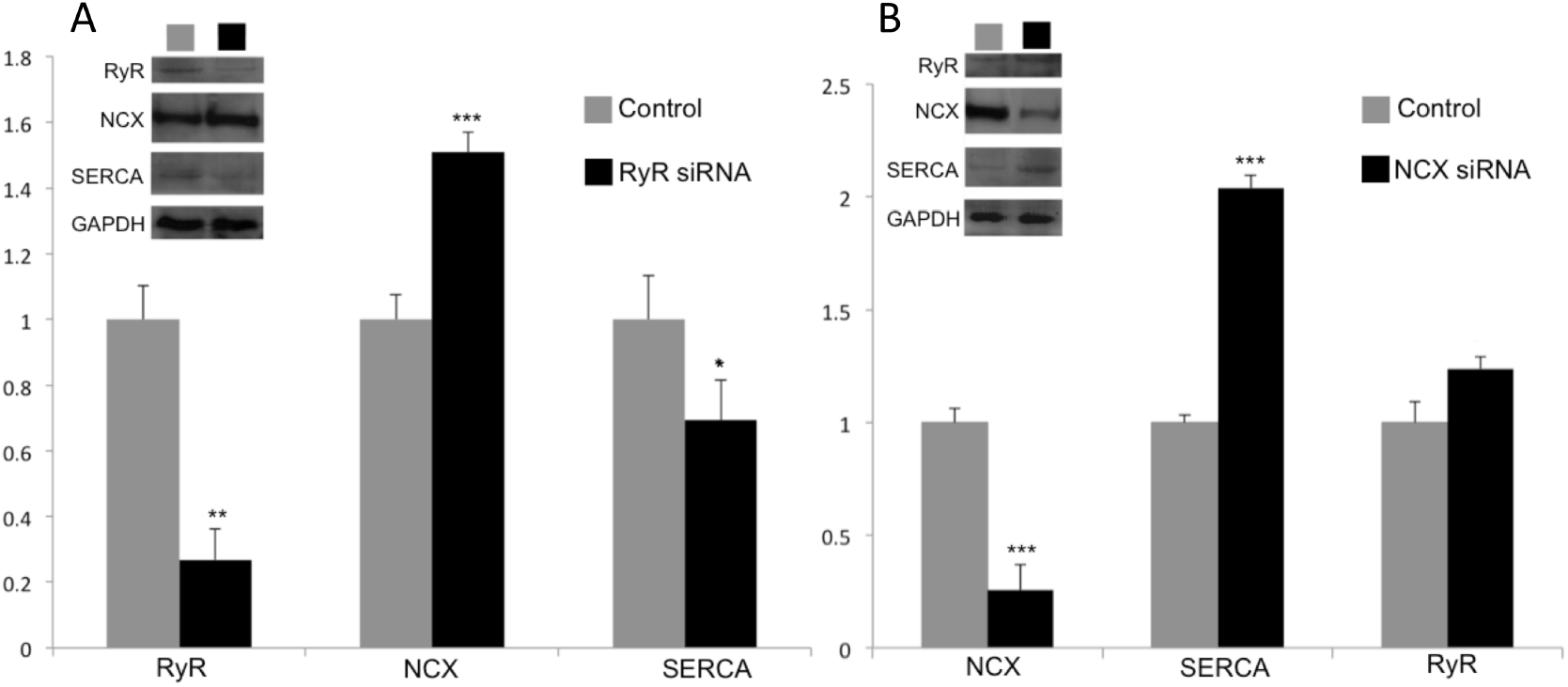
Western blots for RyR knockdown (A) and NCX knockdown (B) efficacy. NRVMs with RyR knockdown experienced increased NCX and decreased SERCA protein expression. NRVMs with NCX knockdown demonstrated increased SERCA expression and statistically insignificant change (P≥0.05) of RyR protein expression.

**FIGURE 2.**
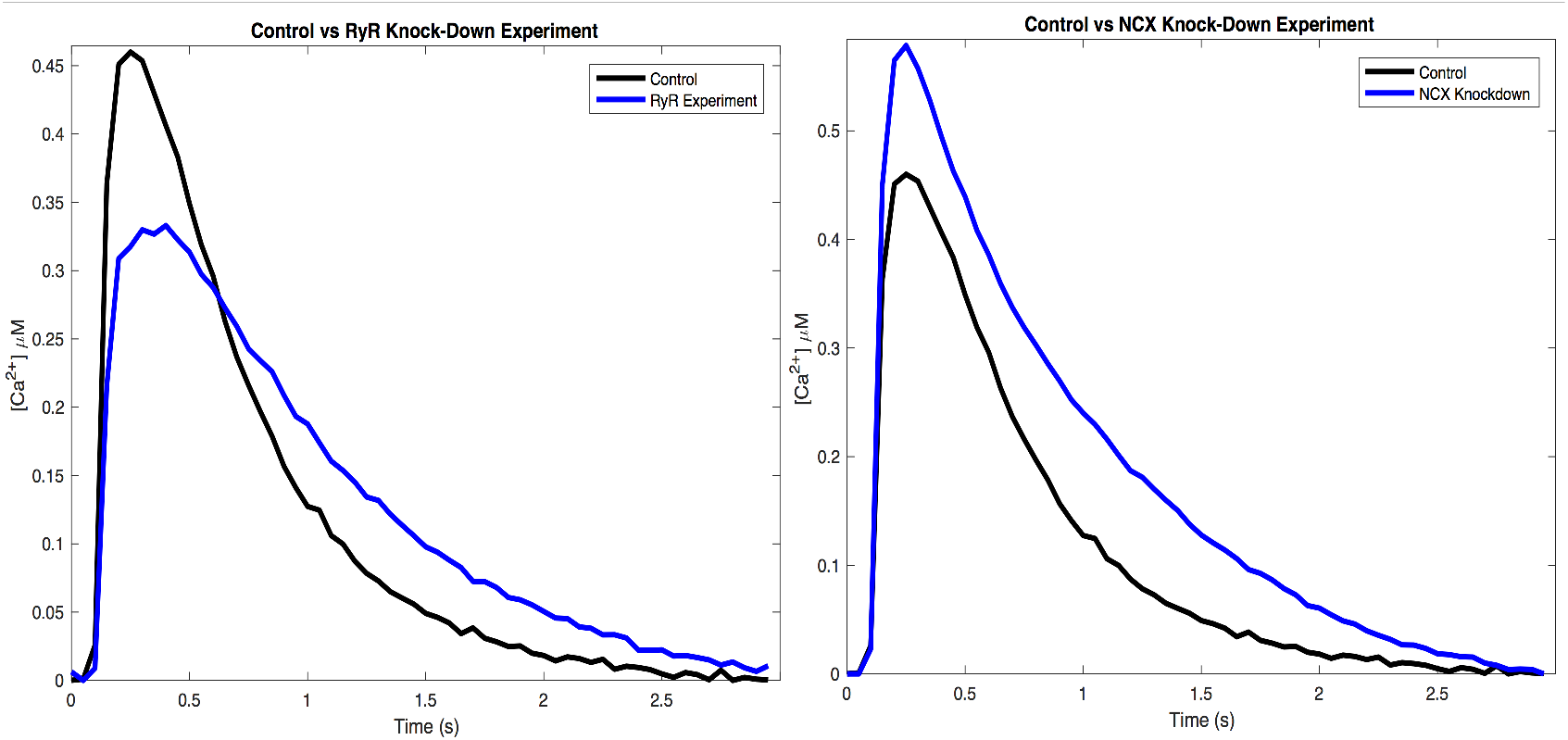
Average Ca^2+^ transients for RyR knockdown (A) and NCX knockdown (B) experimental group with respect to the respective control.

Inhibition of NCX by siRNA showed a decrease of NCX protein expression by 75% (±16% SEM), no statistically significant change in RyR expression, and an increase in SERCA expression by 104%(±2% SEM) (Figure 1B). The associated Ca^2+^ transients showed increased amplitude, and no change to T_peak_ and T_50_ with respect to the control (Figure 2B, Table 1). Similar trends in increased or decreased transcripts of the target protein were observed by qRT-PCR analysis. Transcripts for RyR downregulation experiments decreased RyR by 65%, increased NCX by 75%, and decreased SERCA by 37%. Transcripts for NCX downregulation experiments decreased NCX by 79%, decreased RyR by 3%, and increased SERCA by 56% (Supplementary table 1). There will be some discrepancies in western blot and PCR measurements as there is not a 1:1 relationship in transcript availability and protein expression.

### 3.2 Comparison to Computational Model

We provide the best-fit results from parameter estimates that minimize the weighted sum of squared errors

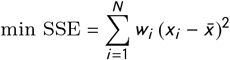

where the weight coefficient is

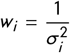

thereby providing experimental data points with less variance, *σ*^2^, greater weight. We perform an initial three-parameter fitting,*k*_*NCX*_, *k*_*R y R*_, and *V*_*max*_, to obtain the right scaling parameters for our control experiments. Parameters used for fitting the data to the model are kept at a minimum to keep the degrees of freedom as small as possible to avoid over-fitting. These parameters are assumed to represent protein expression levels for our control cells (Figure 3, Table 3). Only when necessary are additional parameters used, and only parameters related to the proteins in question are included.

**TABLE 3.** Parameter fitting results

| Parameters | Units | Control | RyR Knockdown | Fold | NCX Knockdown | Fold |
| --- | --- | --- | --- | --- | --- | --- |
| $k_{NCX}$ | $\text{pA}/(\text{pF}(\mu\text{M})^4)$ | $5.08 \times 10^{-16}$ | $7.37 \times 10^{-16}$ | 1.45× | $1.52 \times 10^{-16}$ | 0.3× |
| $d_{NCX}$ | $(\mu\text{M})^4$ | $1 \times 10^{-16}$ * | $5.5 \times 10^{-16}$ | 5.5× | $2 \times 10^{-17}$ | 0.2× |
| $k_{RyR}$ | $\text{ms}^{-1}$ | 0.14 | .041 | 0.3× | 0.14 | 1× |
| $K_{m,RyR}$ | $\mu\text{M}$ | 0.16 * | 0.12 | 0.3× | 0.2 | 1.25× |
| $V_{max}$ | $\mu\text{M}/\text{ms}$ | 0.32 | 0.26 | 0.8× | 0.66 | 2.06× |

Fitting results for the parameters of interest for. Fitted values are reported for the control. RyR and NCX knockdown experiment fitting results are reported. Additionally, fold factors are provided. \* Values are unchanged from [8]

**FIGURE 3.**
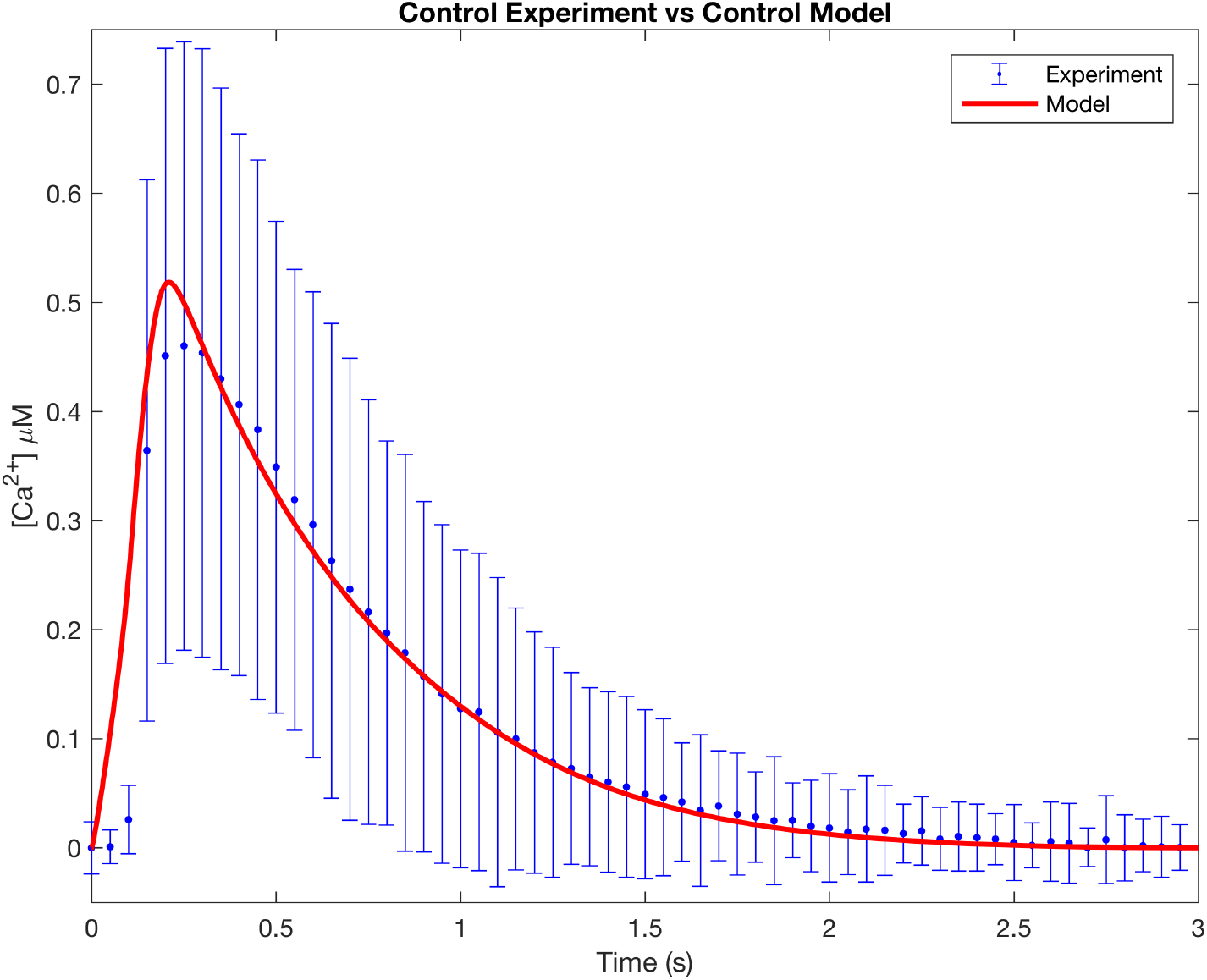
Fitting results for establishing a computational NRVM model for the control experiment. RyR knockdown and NCX knockdown transients were fitted to results of the control model fitting (n = 32).

In order to understand how Ca^2+^ transients respond to the down regulation of key regulatory proteins, we fit the predicted Ca^2+^ transients to the experimental data. For RyR knockdown, we start with only k_*R y R*_ as the fitting parameter, restricted within a range based on the western blot analysis for RyR expression. This limits the parameter search to only within the fractional change corresponding to the western blot data. The model fails to accurately predict experimental data (Figure 4A). Adding the scaling parameters *k*_*NCX*_ for NCX and *V*_*max*_ for SERCA also fail to accurately represent the observed measurements (Figure 4B). Only with the inclusion of the additional parameters *K*_*m, RyR*_ for RyR and *d*_*NCX*_ for NCX do we have a model that is able to predict the experimental Ca^2+^ transient for NRVMs with downregulated RyR (Figure 4C, Table 3).

**FIGURE 4.**
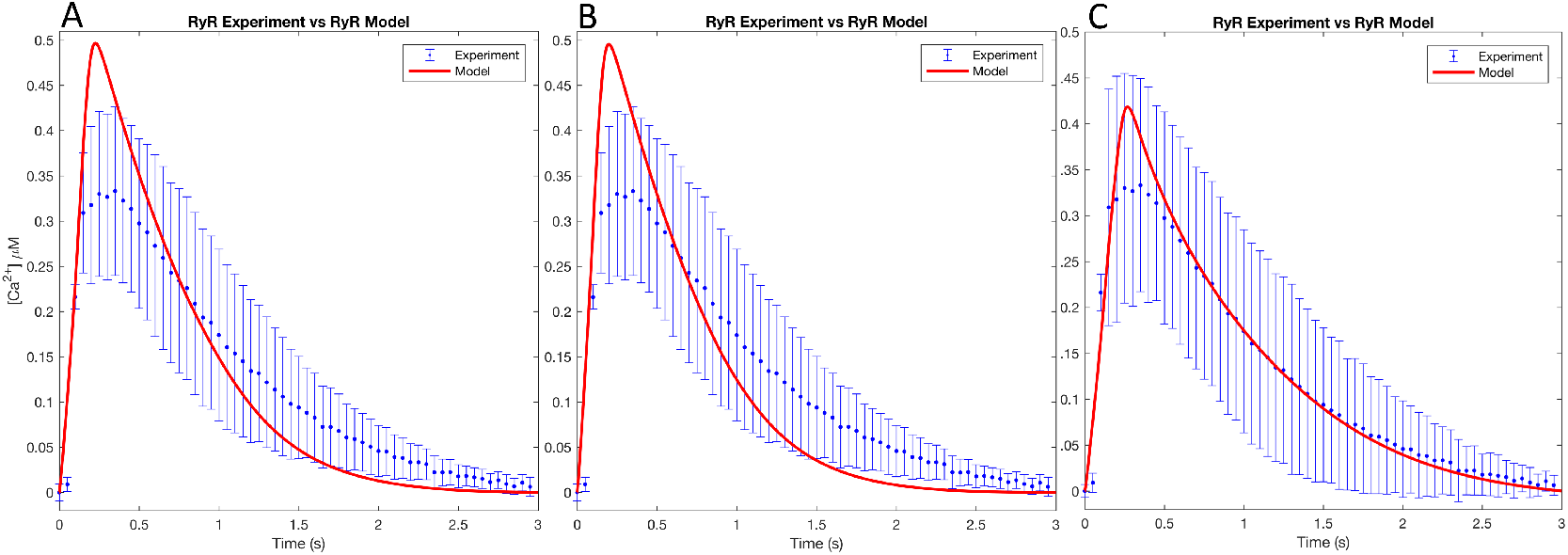
Fitting results using *k*_*R y R*_ as the only parameter (A) did not predict the observed transient. Including additional scaling parameters *k*_*NCX*_ and *v*_*max*_ (B) also failed to predict the experimental transient. Only when *K*_*m, RyR*_ and *d*_*NCX*_ (C) were included with the previous parameter fits did we obtain a representative RyR knockdown Ca^2+^ transient (n = 21).

Similarly, we fit the NCX knockdown experimental Ca^2+^ transients using only the NCX *k*_*NCX*_ parameter. Again, we see that a single parameter fit does not predict the expected Ca^2+^ response (Figure 5A). Fitting with the inclusion of the additional scaling parameters *k*_*R y R*_ and *V*_*max*_ does not accurately predict the NCX knockdown Ca^2+^ transient (Figure 5B). Only by including the additional activity parameters *K*_*m, RyR*_ for RyR and *d*_*NCX*_ for NCX we obtain an accurate prediction of the experimental NCX knockdown Ca^2+^ transient (Figure 5C, Table 3).

**FIGURE 5.**
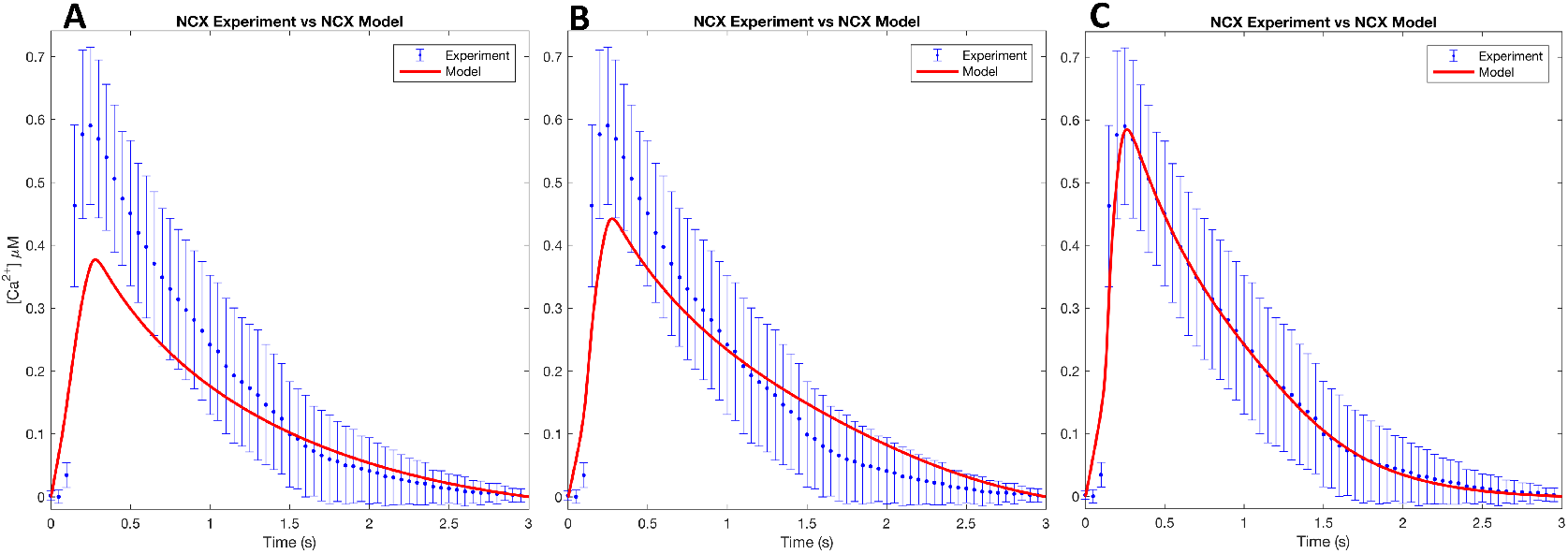
Fitting results using *k*_*NCX*_ as the only parameter (A) did not predict the observed transient. Including additional scaling parameters *k*_*R y R*_ and *v*_*max*_ (B) also failed to predict the experimental transient. Only when *d*_*NCX*_ and *K*_*m, RyR*_ (C) were included with the previous parameter fits did we obtain a representative NCX knockdown Ca^2+^ transient (n = 27).

## 4 DISCUSSION

We present an effort to couple protein expression from NRVMs, their respective [Ca^2+^] transients, and computational modeling to predict changes in cytosolic Ca^2+^ fluxes in the isolated cultured NRVMs when either RyR or NCX are down-regulated by siRNA. While direct comparisons to adult ventricular myocytes is not appropriate, we can compare specific mechanisms in the ECC and CICR process and interpret computational model predictions to generate mechanistic implications. Animal and human studies have shown that protein expression levels of RyR, NCX, and SERCA are altered in diseased states [29, 30] while other experiments with NCX1 knockout mouse models observe no compensatory changes in efflux mechanisms[31]. Furthermore, murine KO models did not show any changes in expression of RyR, SERCA, or Ca^2+^ transients [32, 33]. Our experiments show that healthy NRVMs subjected to downregulation of RyR or NCX demonstrate similar trends in protein expression and Ca^2+^ response as observed in CVD [34]. While insightful, relative expression of Ca^2+^ handling proteins in CVD gives us a snapshot of the current state of the protein levels without mechanistic understanding as to how they reached those levels. Here, we try to provide some answers as to how selective downregulation affects other complimentary proteins and overall regulatory mechanisms of cytoslic [Ca^2+^].

### RyR Downregulation

NRVMs with downregulated RyR have decreased Ca^2+^ amplitude, increased T_peak_, and increase T_50_, suggesting that less Ca^2+^ is being released due to fewer RyR channels and less SR Ca^2+^ store sequestration. Reduction of SERCA expression appears to be a response of SR Ca^2+^ content [35, 36, 37]. Studies have shown lower Ca^2+^ transients and SR store depletion are characteristics of heart failure (HF) arising from depressed SERCA expression and increased NCX expression [38, 39, 40] Increased NCX expression has been reported as compensatory mechanisms to remove cytosolic Ca^2+^ in the presence of reduced SERCA [41, 30, 42]. These differential levels of NCX and SERCA expression in response to siRNA induced RyR knockdown match our experimental observations (Figure 1).

However, the computational model suggests that accounting for RyR, NCX, and SERCA expression levels alone does not recreate the experimental Ca^2+^ transient. Additional fit parameters for RyR such as *k*_*m, RyR*_ for RyR and *d*_*NCX*_ for NCX associated with protein activity are required to accurately predict the experimental observations.

The changes in these additional parameters could be explained by posttranslational modifications of RyR and NCX. *k*_*m, RyR*_ is decreased in the RyR model. A decrease in *k*_*m, RyR*_ implies an increase in RyR activity, possibly from changes in phosphorylation status [43]. This increase in RyR activity could be a response to compensate lower RyR expression. Hyperphosphorylation has been observed in heart failure manifesting as decreased SR Ca^2+^, decreased Ca^2+^ transient amplitude, and increased diastolic [Ca^2+^] possibly from leaky RyRs [40].

The NCX parameter *d*_*NCX*_ is larger in the RyR knockdown model than in the control. An increase in *d*_*NCX*_ corresponds to an increase in NCX activity, which along with an increase in the NCX expression would modulate the cytosolic Ca^2+^ by dampening the transient amplitude [44].

### NCX Downregulation

NRVMs with downregulated NCX have increased Ca^2+^ amplitudes while maintaining T_peak_ and T_50_. Increases in Ca^2+^ SR storage have been reported when NCX efflux is decreased. SERCA upregulation compensates for the lack of cytosolic Ca^2+^ extrusion by NCX. The increase in SR Ca^2+^ results in increased systolic Ca transients [45]. The upregulation of SERCA in response to NCX knockdown is observed in our experiments (Figure 1). However, once again accounting for only the protein expression levels does not adequately predict the Ca^2+^ transient (Figure 5A and 5B). However, when we include the additional NCX and RyR parameters, we do have an accurate prediction of the expected Ca^2+^ response.

As mentioned above, NCX’s *d*_*NCX*_ parameter serves to modulate the Ca^2+^ transient. Decreased NCX expression does not account for the increase in Ca^2+^ amplitude. NCX needs to be less active, decreasing Ca^2+^ efflux. This can be explained by the decrease in *d*_*NCX*_. Similarly, RyR has to compensate for an increase in SR content. We see an increase *k*_*m, RyR*_ with RyR being less sensitive to Ca^2+^, possibly to compensate for increase cytosolic [Ca^2+^]. Decrease in RyR activity may be attributed to decreased phosphorylation due to increased luminal Ca^2+^. Luminal SR [Ca^2+^] is known to have a close interrelation between SR function and SR [Ca^2+^] which affects Ca^2+^ binding, Ca^2+^ release channels, and Ca^2+^ pumps [46].

## 5 CONCLUSION

Ca^2+^ handling is a function of developmental stage and disease state [4, 24, 23, 47, 48]. Expression levels of NCX and SERCA in rat cardiocytes alter significantly with development. Post-natal rats express high NCX and low SERCA levels while adult rats have low NCX and high SERCA levels [49]. As expected, differences is cytosolic Ca^2+^ transient decline due to NCX and SERCA are observed in adult vs. neonate rats. In adult rats, Ca^2+^ removal by NCX accounts for 7% while SERCA removes 92%. Contrasted with neonates, NCX removes 46% and SERCA removes 50% [50]. Similarly, murine models demonstrate drastically significant integrated Ca^2+^ flux handling with adult cardiocytes removing 91% of calcium to the SR and 3% removed by NCX while neonates remove 70% to the SR and 24% extruded by NCX [51]. With calcium flux varying between developmental stages, we consider comparisons between NCX1 KO models in adult myocytes and downregulation in NRVMs. Does an 80-90% NCX1 KO, responsible for 3% of calcium removal impact calcium flux in a manner observed by 70-80% NCX downregulation in NRVMs? Answering these questions would expand our understanding of cardiac disease as a congenital or acquired condition with respect to developmental stage.

The experimental and computational results presented here show that protein expressions of key Ca^2+^ regulatory channels are interdependent. Additionally, the activity of these channels is co-regulated to compensate for changes in the Ca^2+^ transient and the cytosolic Ca^2+^ concentration. Our understanding of how Ca^2+^ cycling proteins respond to induced downregulation can help us create a response pathway in posttranslational mechanisms to generate and test new hypothesis. With Ca^2+^ involvement in so many signaling processes, experiments identifying which pathways are activated when RyR or NCX are downregulated are needed. This work is limited by the number of proteins measured and use of cell-wide calcium measurement. Measuring if any change in LTCC Ca^2+^ flux and SR content would further help validate claims that the observed changes in Ca^2+^ transient amplitude are due to increased or decreased SR Ca^2+^ storage. Proteomic and genomic studies could be used quantify how accessory proteins adapt to induced protein inhibition and identify which pathways are actively responsible for changes in protein expression.

