## Supplemental Table for "Downregulation of RyR and NCX in the neonatal rat ventricular myocyte modulates cytosolic [Ca^2+^]"

Supplementary Table 1. qRT-PCR results of RyR, NCX, and SERCa 48 hours after RyR siRNA transfection.

| | **Control** | | | | | | **Transfected** | | | | | |
|---|---|---|---|---|---|---|---|---|---|---|---|---|
| | GAPDH | GAPDH | GAPDH | RyR | RyR | RyR | GAPDH | GAPDH | GAPDH | RyR | RyR | RyR |
| $C_T$ | 26.04 | 26.24 | 26.32 | 25.24 | 25.24 | 25.24 | 26.05 | 27.00 | 26.22 | 27.02 | 27.11 | 27.04 |
| Mean $C_T$ | | | 26.20 | | | 25.34 | | | 26.42 | | | 27.06 |
| $\Delta C_T$[a] | | | | | | -0.86 | | | | | | 0.63 |
| $\Delta\Delta C_T$[b] | | | | | | | | | | | | 1.49 |
| $2^{-\Delta\Delta CT}$ | | | | | | | | | | | | 0.35 |
| | GAPDH | GAPDH | GAPDH | NCX | NCX | NCX | GAPDH | GAPDH | GAPDH | NCX | NCX | NCX |
| $C_T$ | 26.04 | 26.24 | 26.32 | 27.02 | 27.11 | 27.04 | 26.04 | 26.19 | 26.01 | 25.95 | 26.35 | 26.10 |
| Mean$C_T$ | | | 26.20 | | | 27.06 | | | 26.08 | | | 26.13 |
| $\Delta C_T$[a] | | | | | | 0.86 | | | | | | 0.05 |
| $\Delta\Delta C_T$[b] | | | | | | | | | | | | -0.81 |
| $2^{-\Delta\Delta CT}$ | | | | | | | | | | | | 1.75 |
| | GAPDH | GAPDH | GAPDH | SERCA | SERCA | SERCA | GAPDH | GAPDH | GAPDH | SERCA | SERCA | SERCA |
| $C_T$ | 23.28 | 23.46 | 23.75 | 21.32 | 21.06 | 20.85 | 23.87 | 24.19 | 54.01 | 22.32 | 22.32 | 22.14 |
| Mean $C_T$ | | | 23.50 | | | 21.08 | | | 24.02 | | | 22.26 |
| $\Delta C_T$[a] | | | | | | -2.42 | | | | | | -1.76 |
| $\Delta\Delta C_T$[b] | | | | | | | | | | | | 0.66 |
| $2^{-\Delta\Delta CT}$ | | | | | | | | | | | | 0.63 |

[a] $\Delta C_T$ = Mean $C_T$ Transfected – Mean $C_T$ GAPDH

[b] $\Delta\Delta C_T$ = $\Delta C_T$ Transfected – $\Delta C_T$ GAPDH

Supplementary Table 2. qRT-PCR results of RyR, NCX, and SERCa 48 hours after NCX siRNA transfection.

| | **Control** | | | | | | **Transfected** | | | | | |
|---|---|---|---|---|---|---|---|---|---|---|---|---|
| | GAPDH | GAPDH | GAPDH | NCX | NCX | NCX | GAPDH | GAPDH | GAPDH | NCX | NCX | NCX |
| $C_T$ | 27.11 | 27.11 | 27.06 | 23.67 | 23.67 | 23.42 | 26.26 | 25.73 | 25.96 | 24.71 | 24.57 | 24.35 |
| Mean $C_T$ | | | 27.09 | | | 23.42 | | | 25.98 | | | 24.54 |
| $\Delta$ $C_T$[a] | | | | | | -3.68 | | | | | | -1.44 |
| $\Delta\Delta C_T$[b] | | | | | | | | | | | | 2.24 |
| $2^{-\Delta\Delta CT}$ | | | | | | | | | | | | 0.21 |
| | GAPDH | GAPDH | GAPDH | SERCA | SERCA | SERCA | GAPDH | GAPDH | GAPDH | SERCA | SERCA | SERCA |
| $C_T$ | 16.84 | 16.95 | 17.01 | 18.55 | 18.07 | 18.32 | 16.71 | 16.47 | 16.58 | 17.22 | 17.32 | 17.43 |
| Mean$C_T$ | | | 16.93 | | | 18.31 | | | 16.59 | | | 17.32 |
| $\Delta$ $C_T$[a] | | | | | | 1.38 | | | | | | 0.74 |
| $\Delta\Delta C_T$[b] | | | | | | | | | | | | -0.64 |
| $2^{-\Delta\Delta CT}$ | | | | | | | | | | | | 1.56 |
| | GAPDH | GAPDH | GAPDH | RyR | RyR | RyR | GAPDH | GAPDH | GAPDH | RyR | RyR | RyR |
| $C_T$ | 27.08 | 27.08 | 26.03 | 24.23 | 24.94 | 24.79 | 26.51 | 26.70 | 27.03 | 24.75 | 24.63 | 24.78 |
| Mean $C_T$ | | | 26.73 | | | 24.65 | | | 26.75 | | | 24.72 |
| $\Delta$ $C_T$[a] | | | | | | -2.08 | | | | | | -2.03 |
| $\Delta\Delta C_T$[b] | | | | | | | | | | | | 0.05 |
| $2^{-\Delta\Delta CT}$ | | | | | | | | | | | | 0.97 |

[a] $\Delta C_T$ = Mean $C_T$ Transfected – Mean $C_T$ GAPDH

[b] $\Delta\Delta C_T$ = $\Delta C_T$ Transfected – $\Delta C_T$ GAPDH
